# *De novo* synthesis of phosphatidylcholine is essential for the promastigote but not amastigote stage in *Leishmania major*

**DOI:** 10.1101/2020.12.30.424847

**Authors:** Samrat Moitra, Somrita Basu, Mattie Pawlowic, Fong-fu Hsu, Kai Zhang

## Abstract

Phosphatidylcholine (PC) is the most abundant type of phospholipids in eukaryotes constituting ~30% of total lipids in *Leishmania*. PC synthesis mainly occurs via the choline branch of the Kennedy pathway (choline ⇒ choline-phosphate ⇒ CDP-choline ⇒ PC) and the N-methylation of phosphatidylethanolamine (PE). In addition, *Leishmania* parasites can acquire PC and other lipids from the host or culture medium. In this study, we assessed the function and essentiality of choline ethanolamine phosphotransferase (CEPT) in *Leishmania major* which is responsible for the final step of the *de novo* synthesis of PC and PE. Our data indicate that CEPT is localized in the endoplasmic reticulum and possesses the activity to generate PC from CDP-choline and diacylglycerol. Targeted deletion of *CEPT* is only possible in the presence of an episomal *CEPT* gene in the promastigote stage of *L. major*. These chromosomal null parasites require the episomal expression of *CEPT* to survive in culture, confirming its essentiality during the promastigote stage. In contrast, during *in vivo* infection of BALB/c mice, these chromosomal null parasites appeared to lose the episomal copy of *CEPT* while maintaining normal levels of virulence, replication and cellular PC. Therefore, while the *de novo* synthesis of PC/PE is indispensable for the proliferation of promastigotes, intracellular amastigotes appear to acquire most of their lipids through salvage and remodeling.

## INTRODUCTION

Leishmaniasis is caused by protozoan parasites of the genus *Leishmania* which alternate between extracellular promastigotes colonizing the midgut of sandflies and intracellular amastigotes residing in the macrophage of mammals (Alvar et al., 2012). Without a safe vaccine, the mitigation of leishmaniasis mainly depends on vector control and drugs (Croft and Olliaro, 2011). Current therapeutics are limited and plagued with high toxicity (Croft and Olliaro, 2011). A better understanding of how *Leishmania* parasites acquire essential cellular components may lead to new drug targets and improved treatments.

Glycerophospholipids such as phosphatidylcholine (PC) and phosphatidylethanolamine (PE) are among the most abundant types of lipids accounting for approximately 30% and 10% of total lipids in *Leishmania*, respectively (Zhang and Beverley, 2010; Zheng et al., 2010). While PC in promastigotes primarily consists of 1,2-diacyl-PC, the majority of leishmanial PE belongs to 1-O-alk-1′-enyl-2-acyl-*sn*-glycero-3-phosphoethanolamine or plasmenylethanolamine (PME) (Beach et al., 1979; Zufferey et al., 2003; Pawlowic et al., 2016; Moitra et al., 2019). Due to its positively charged head group, PC is a membrane-forming phospholipid that is more abundant on the outer leaflet of the plasma membrane (van Meer et al., 2008; Nickels et al., 2015). Meanwhile, PE is known for its ability to promote membrane fusion and mainly resides in the inner leaflet (Verkleij et al., 1984; Ellens et al., 1989; Zachowski, 1993). Besides being principal membrane components, PC and PE possess other important functions. For example, PC can serve as the precursor for a number of important signaling molecules and metabolic intermediates including lyso-PC, phosphatidic acid, diacylglycerol (DAG) and free fatty acids (FA) (Exton, 1994; Furse and de Kroon, 2015). In addition, PE is required for the synthesis of GPI-anchored proteins in trypanosomatids by providing the ethanolamine (EtN) phosphate bridge that links proteins to glycan anchors (Menon et al., 1993). PE also contributes to the formation of autophagosome during differentiation and starvation (Ichimura et al., 2000; Besteiro et al., 2006; Williams et al., 2012) and the posttranslational modification of eukaryotic elongation factor 1A in trypanosomatids (Signorell et al., 2008a). Thus, understanding the mechanism by which *Leishmania* acquire their PC and PE may reveal new ways to block their growth.

In many eukaryotic cells, the majority of PE and PC are synthesized *de novo* via the Kennedy pathway (Kennedy, 1956; Cui and Vance, 1996; Gibellini and Smith, 2010). As indicated in Fig. 1, the *de novo* synthesis of PE, i.e. the EtN branch of the Kennedy pathway, starts with the phosphorylation of EtN into EtN-phosphate, which is also generated from the metabolism of sphingoid bases (Zhang et al., 2007); EtN-phosphate is then conjugated to CTP by ethanolaminephosphate cytidylyltransferase (EPCT) to produce CDP-EtN and pyrophosphate; and two enzymes catalyze the final steps: an ethanolamine phosphotransferase (EPT) which combines CDP-EtN with 1-alkyl-2-acyl-glycerol to form the precursor of PME (Pawlowic et al., 2016) (Fig. 1), and a choline ethanolamine phosphotransferase (CEPT) which condenses CDP-EtN and DAG into diacyl-PE (Fig. 1). A parallel route, aka the choline branch of the Kennedy pathway, is responsible for the *de novo* synthesis of PC (choline → choline-phosphate → CDP-choline → PC, Fig. 1), and CEPT catalyzes the last step of combining CDP-choline and DAG into PC as a dual activity enzyme (Henneberry and McMaster, 1999; Signorell et al., 2008b; Vance, 2008).

**Figure 1.**
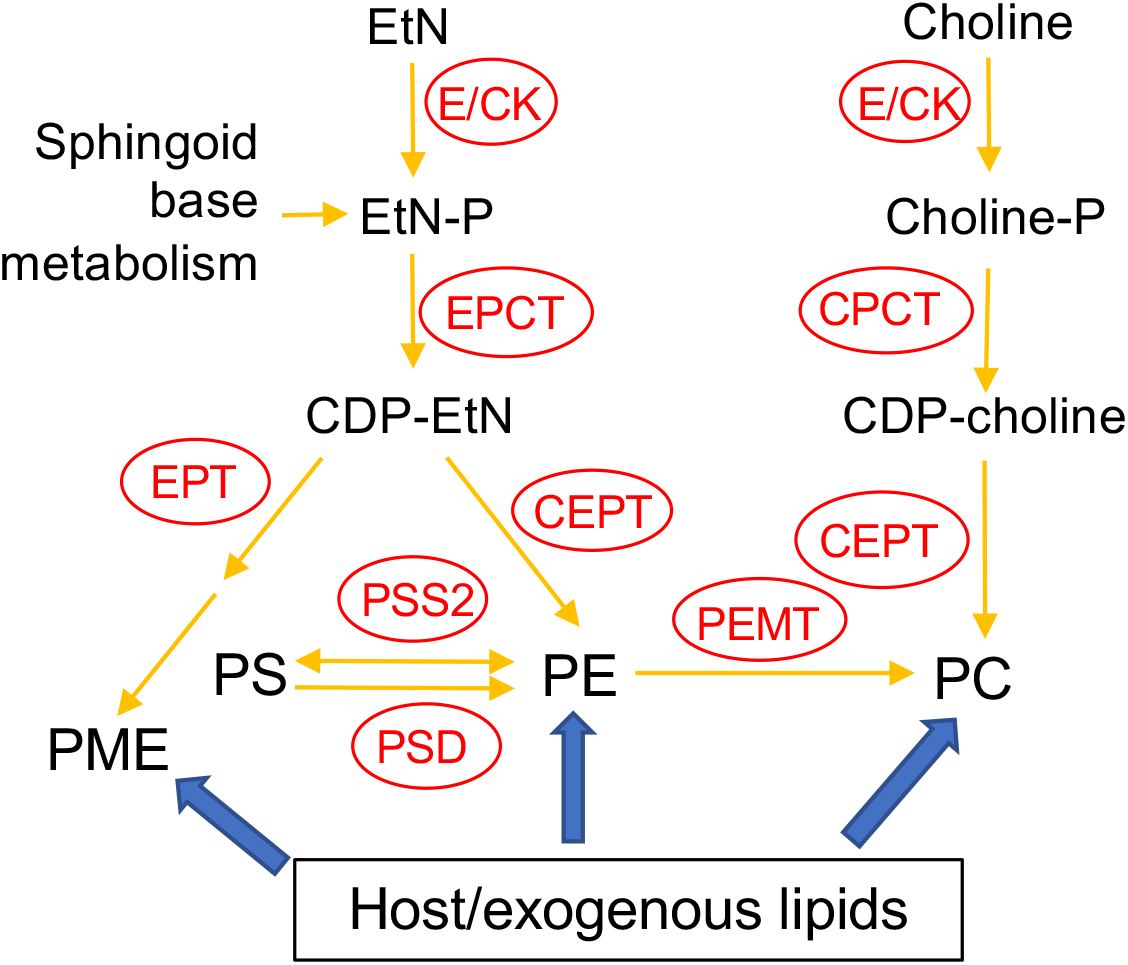
Acquisition of PE and PC in *Leishmania*. E/CK: ethanolamine/choline kinase; EPCT: ethanolaminephosphate cytidylyltransferase; EPT: ethanolamine phosphotransferase; CPCT: cholinephosphate cytidylyltransferase; CEPT: choline ethanolamine phosphotransferase; PEMT: phosphatidylethanolamine N-methyltransferase; PSS2: phosphatidylserine synthase 2; PSD: phosphatidylserine decarboxylase; EtN: ethanolamine; EtN-P: ethanolamine phosphate; PE: (1,2-diacyl-) phosphatidylethanolamine; PME: plasmenylethanolamine; Choline-P: choline phosphate; PC: phosphatidylcholine; PS: phosphatidylserine.

In addition to *de novo* synthesis, PE and PC are also generated from the conversion of phosphatidylserine (PS) and the N-methylation of PE, respectively (Fig. 1). Furthermore, *Leishmania* can scavenge glycerophospholipids and degrade them via the activity of phospholipase A2 (PLA2), suggesting that these parasites can remodel exogenous lipids into their own via the Lands cycle (Das et al., 2001; Henriques et al., 2003; Parodi-Talice et al., 2003; Castanys-Munoz et al., 2007; Pawlowic and Zhang, 2012) (Fig. 1). Because the Kennedy pathway is the dominant route for PE/PC synthesis in most eukaryotes including *Trypanosoma brucei* (a trypanosomatid parasite related to *Leishmania*) (Signorell et al., 2008b; Gibellini and Smith, 2010), we have examined the roles of several genes involved in this pathway in *Leishmania major*. First, to explore the significance of PME production, we generated the *EPT*-null mutants (*ept*^*−*^) which were largely devoid of PME but contained WT-levels of diacyl-PE (Pawlowic et al., 2016). *Ept*^*−*^ promastigotes were fully viable and replicative in culture but showed significantly attenuated virulence in mice (Pawlowic et al., 2016). Meanwhile, *ept*^*−*^ amastigotes were fully virulent, indicating that PME synthesis alone was not required during the mammalian stage for *L. major* (Pawlowic et al., 2016). Second, to determine the role of *de novo* PC synthesis in *L. major*, we focused on the cholinephosphate cytidylyltransferase (CPCT) which catalyzes the choline-phosphate to CDP-choline conversion in the choline branch of the Kennedy pathway (Fig. 1). Without CPCT, *L. major* parasites could not incorporate choline into PC, yet the *CPCT*-null promastigotes (*cpct*^*−*^) contained similar levels of PC and PE as WT parasites in culture (Moitra et al., 2019). Loss of *CPCT* did not affect promastigote replication in complete media but caused reduced growth rates under EtN-limiting conditions (Moitra et al., 2019). Both *cpct*^*−*^ promastigotes and amastigotes were fully virulent in mice (Moitra et al., 2019). Collectively, these observations suggest that other routes of PC synthesis (e.g., PE N-methylation and lipid salvage) can compensate the loss of CPCT (Fig. 1).

In this study, we investigated the function of CEPT in *L. major* which is expected to directly catalyze the formation of diacyl-PC and diacyl-PE via the Kennedy pathway (Fig. 1). In *T. brucei*, RNAi knockdown of CEPT significantly reduced the biosynthesis of PE and PC and caused a cytokinesis defect resulting in the accumulation of zoids (Signorell et al., 2008b; Signorell et al., 2009). These findings are in agreement with the dual substrate activity exhibited by *T. brucei* CEPT (Farine et al., 2015). Given the fact that CPCT is not essential for *L. major*, it is important to probe whether the *de novo* PC synthesis is required for *Leishmania* promastigotes and amastigotes. Unlike *T. brucei* which has separate Kennedy pathway branches for PE and PC synthesis (Gibellini and Smith, 2010; Smith and Butikofer, 2010), *Leishmania* parasites can incorporate EtN into both PE and PC (Pawlowic et al., 2016; Moitra et al., 2019), presumably through the activity of PE-N-methyltransferases (Bibis et al., 2014). Thus, CEPT is likely required for the synthesis of PC through both the Kennedy pathway and the methylation of PE in *Leishmania* (Fig. 1). Our results demonstrate that CEPT is indispensable for *L. major* promastigotes but not amastigotes, suggesting that intracellular parasites can fulfill their need for PC through lipid salvage/remodeling.

## MATERIALS AND METHODS

### Materials

For the CEPT activity assay, [methyl-^14^C] CDP-choline (50–60 mCi/mmol) and [diarachidonyl-1-^14^C] PC (50–60 mCi/mmol) were purchased from Perkin Elmer, Inc (Waltham, MA) and American Radiolabeled Chemicals (St. Louis, MO), respectively. 1,2-Dioctanoyl-sn-glycerol (DAG) was purchased from EMD Millipore (Burlington, MA). Lipid standards for mass spectrometry including 1,2-dimyristoyl-*sn*-glycero-3-phosphoethanolamine (14:0/14:0-PE) and 1,2-dimyristoyl-*sn*-glycerol-3-phosphocholine (14:0/14:0-PC) were purchased from Avanti Polar Lipids (Alabaster, AL). All other reagents were purchased from Thermo Fisher Scientific (Hampton, NH) unless specified otherwise.

### Molecular constructs

The open reading frame (ORF) of *CEPT* (LmjF36.5900) was amplified by PCR from *L. major* genomic DNA and cloned into the pXG vector (Ha et al., 1996) to generate pXG-*CEPT*. The *CEPT* ORF was then subcloned into the pXG-GFP’ (Ha et al., 1996) and pXNG4 vectors (Murta et al., 2009) to generate pXG-*GFP-CEPT* and pXNG4-*CEPT*, respectively. The upstream and downstream flanking sequences (~1 Kb each) of *CEPT* were PCR amplified and cloned in tandem into the pUC18 vector. Genes conferring resistance to puromycin (*PAC*) and blasticidin (*BSD*) were inserted between the upstream and downstream flanking sequences to generate pUC18-KO-*CEPT:PAC* and pUC18-KO-*CEPT:BSD*, respectively. All the molecular constructs were confirmed by restriction enzyme digestion and sequencing. Oligo nucleotides used in this study are summarized in Table S2.

### *Leishmania* culture and genetic manipulation

*L. major* LV39 clone 5 (Rho/Su/59/P) promastigotes were cultivated at 27 °C in M199 media with 10% heat-inactivated fetal bovine serum and other supplements as previously described (Kapler et al., 1990). To monitor cell growth, promastigotes were inoculated at 1.0-2.0 × 10^5^ cells/ml and culture densities were determined over time using a hemocytometer. In general, log phase promastigotes refer to replicative parasites at densities lower than 1.0 × 10^7^ cells/ml and stationary phase promastigotes refer to mainly non-replicative parasites at 2.0-3.0 × 10^7^ cells/ml. The proliferation of promastigotes from 1.0-2.0 × 10^5^ cells/ml to 2.0-3.0 × 10^7^ cells/ml (usually in 2-3 days) is referred to as one passage.

To delete *CEPT*, antibiotic resistance cassettes acquired from pUC18-KO-*CEPT:PAC* and pUC18-KO-*CEPT:BSD* were sequentially introduced into LV39 wild type (WT) parasites and drug selections were performed as previously described (Kapler et al., 1990). With this approach, we were able to generate the heterozygous *CEPT+/−* mutants *(ΔCEPT::PAC/CEPT*) but not the homozygous *cept* ^*−*^ mutants (*ΔCEPT::PAC/ΔCEPT::BSD*). To delete the second chromosomal allele of *CEPT*, the pXNG4-*CEPT* plasmid containing *GFP*, *TK*, *SAT* (nourseothricin resistance) and *CEPT* genes was introduced into *CEPT+/−* parasites. The resulting *CEPT+/−* +pXNG4-*CEPT* cells were transfected with the blasticidin resistance cassette from pUC18-*CEPT-KO:BSD*. Parasite showing resistance to puromycin, blasticidin, and nourseothricin were serially diluted into 96-well plates. The resulting clones were amplified and processed for Southern blot analysis to verify the loss of chromosomal *CEPT* and referred to as *cept* ^*−*^ +pXNG4-*CEPT*.

### Fluorescence microscopy and validation of GFP-*CEPT*

Promastigotes expressing GFP-*CEPT* were attached to poly-L-lysine coated coverslips, fixed with 3.7% formaldehyde, and then permeabilized on ice with ethanol. Incubation with rabbit anti-*T. brucei* BiP antiserum (1:1000) was performed at room temperature for 40 min. After washing three times with phosphate-buffered saline (PBS), coverslips were incubated with a goat anti-rabbit-IgG Texas Red (1:2000) antiserum for 40 min. After washing three times with PBS, DNA was stained with 2.5 μg/ml of Hoechst 33342, followed by final three washes with PBS. Coverslips containing cells were mounted on glass slides with Fluormount-G mounting medium and an Olympus BX51 Upright Fluorescence Microscope was used to visualize the expression and localization of GFP-*CEPT*. To quantify the overlap between GFP-*CEPT* and anti-BiP staining, 136 randomly selected cells were analyzed using the Image J JACoP (Just Another Colocalization Plugin) (Table S1) (Bolte and Cordelieres, 2006).

To detect GFP-*CEPT* by Western blot, *Leishmania* promastigotes were boiled in SDS sample buffer, resolved by SDS-PAGE and probed with a rabbit anti-GFP polyclonal antibody (Life Technologies) followed by a goat anti-rabbit IgG-HRP antibody. To ensure equal loading, blots were probes with an anti-α-tubulin monoclonal antibody followed by a goat anti-mouse IgG-HRP antibody. Signals from Western blot were detected using a FluorChem E system (Protein Simple).

### CEPT assay and thin layer chromatography (TLC)

To examine the CEPT activity in *Leishmania*, log phase promastigotes of WT, *CEPT+/− +*pXG-*CEPT* and WT *+*pXG-*GFP-CEPT* were harvested by centrifugation, washed twice with PBS, and resuspended in a hypotonic lysis buffer (1 mM of Tris-HCl, 0.1 mM of EDTA, 1 × protease inhibitor cocktail, pH 8) at 5 × 10^8^ cells/ml. Cells were lysed by sonication (Ultrasonic homogenizer Sonic Ruptor 250, medium speed, 30 s × 3 times on ice) and protein concentration in lysate was determined by the BCA assay. Procedure for the CEPT activity assay was adapted from a previously reported method with *Trypanosoma brucei* cells (Farine et al., 2015). Briefly, each 100-μl reaction contained 900 μg of leishmanial protein (from promastigote lysate), 50 μM of [^14^C] CDP-choline, 50 μM of DAG, 50 mM of Tris pH 8.0, 10 mM of MgCl2, and 0.005% of Tween 20 (w/v). After 1-hour incubation at room temperature, the reaction mix was extracted twice with of 250 μl of 1-butanol. The combined organic fractions were dried under a nitrogen stream and reconstituted in 20 μl of 1-butanol. The aqueous fraction was also saved. Materials from the organic and aqueous fractions were analyzed by one-dimensional TLC on a silica gel 60 plate (Merck) using a solvent system composed of chloroform:methanol:acetate:water (25:15:4:2 by volumes). [^14^C] CDP-choline, [^14^C] 1,2-diarachidonyl PC, and [^14^C] choline chloride were used in the TLC as markers. Mouse liver homogenate was used as a positive control and boiled *Leishmania* lysates were used as negative controls. After TLC, radioactive signals on the silica gel 60 plate were detected using a Personal Molecular Imager (Bio-Rad).

### GCV treatment of promastigotes in culture

Promastigotes of WT, *CEPT+/−* +pXNG4-*CEPT* and *cept* ^*−*^ +pXNG4-*CEPT* were cultivated in complete M199 media in the absence or presence of 50 μg/ml of GCV or 200 μg/ml of nourseothricin (for *CEPT+/−* +pXNG4-*CEPT* and *cept* ^*−*^ +pXNG4-*CEPT* only). Every three days, parasites were re-inoculated at 1.0 × 10^5^ cells/ml in fresh media containing the same concentration of GCV or nourseothricin to start a new passage. For every passage, the GFP expression level was determined by flow cytometry at mid-log phase (3-7 × 10^6^ cells/ml) using an Attune NxT Acoustic Flow Cytometer. After 16 passages in the presence of 50 μg/ml of GCV, *cept*^*−*^ *+*pXNG4-*CEPT* parasites were sorted based on their GFP expression levels using a BD Biosciences FACS Aria III Plus Cell Sorter. GFP-high and GFP-low parasites (~50,000 each) were sorted directly into complete M199. Parasites showing low GFP expression were serially diluted into 96-well plates. Multiple clones were then selected and their GFP expression levels were determined by flow cytometry (examples are shown in Fig. 5E-G).

### Quantitative PCR (qPCR) analyses to determine *CEPT* transcript level, parasite loads and plasmid copy numbers

To determine the *CEPT* transcript level in WT parasites, total RNA was extracted from log phase promastigotes or lesion-derived amastigotes. 1 μg of RNA was converted into complementary DNA using a high-capacity cDNA conversion kit, followed by qPCR using primers designed for the ORF of *CEPT* or the 28S rRNA gene. The expression level of *CEPT* was normalized to that of 28S rRNA using the comparative Ct approach, also known as the 2^−ΔΔ (Ct)^ method (Livak and Schmittgen, 2001).

To determine parasite numbers, a standard curve was prepared from the serially diluted genomic DNA samples extracted from WT promastigotes (from 2 × 10^6^ cells to 0.2 cells per reaction). Dilutions were carried out in the presence of salmon sperm DNA as a carrier. QPCR was performed on DNA extracted from promastigotes and amastigotes with primers for the 28S rRNA gene. Parasite numbers were determined using the standard curve.

To determine pXNG4-*CEPT* copy numbers, a calibration curve was generated by serially diluting pXNG4-*CEPT* plasmid DNA from 2 × 10^6^ molecules to 0.2 molecules per reaction in the presence of salmon sperm DNA. QPCR was performed on DNA extracted from promastigotes and amastigotes using primers for the pXNG4 plasmid. Plasmid copy number per cell was calculated by dividing the total plasmid copy number by the total parasite number. All qPCR reactions were performed in triplicates using the SSO Advanced Universal SYBR Green Supermix (Bio-Rad) in an Applied Biosystems 7300 RT-PCR thermo-cycler.

### Mouse footpad infection and in vivo GCV treatment

Use of mice in this study was approved by the Animal Care and Use Committee at Texas Tech University. BALB/c mice (female, 8 weeks old) were purchased from Charles River Laboratories International. To assess parasite virulence, stationary phase promastigotes were resuspended in serum free M199 medium and injected into the left hind footpad of BALB/c mice (1.0 × 10^6^ cells/mouse, 10 mice per group). For each group, one half of the mice received GCV treatment at 7.5 mg/kg/day (GCV was prepared in sterile PBS, 0.5 ml per day) through intraperitoneal injection, while the other half received sterile PBS (0.5 ml per day). GCV and PBS treatments started one day post infection and continued for 14 days. Mouse body weights were monitored once a week for three weeks post injection. Footpad lesions were measured weekly using a Vernier caliper. Parasite loads in infected footpads were determined at the indicated times by limiting dilution assay as described previously (Titus et al., 1985) and qPCR as described above. Mice were euthanized via CO_2_ asphyxiation when their footpad lesions exceeded 2.5 mm or when secondary infections were detected.

### Recovery and analyses of promastigotes from infected mice

Amastigotes were isolated from 8-10 weeks infected mice and allowed to convert into promastigotes in complete M199 media in the presence or absence of 200 μg/ml of nourseothricin or 50 μg/ml of GCV. Parasites were then continuously cultivated in media containing the same concentration of nourseothricin or GCV. During each passage, the growth rate and GFP expression were determined by cell counting and flow cytometry as described above.

### Lipid analysis by electrospray ionization mass spectrometry (ESI-MS)

Total lipids from log phase promastigotes, lesion-derived amastigotes, or uninfected mouse tissue were extracted using the Bligh-Dyer method (Bligh and Dyer, 1959; Zhang, 2012). Purification of lesion amastigotes was performed as previously described (Glaser et al., 1990; Zhang, 2012). Identification of PC, PE and TAG structures using low-energy collision induced dissociation LIT MS^n^ with high-resolution Fourier transform mass spectrometry was conducted on a Thermo Scientific LTQ Orbitrap Velos mass spectrometer (R=100,000 at *m/z* 400) with Xcalibur operating system as previously described (Hsu et al., 2014). Fatty acyl constituents and numbers of double bonds in PC, PE and TAG were given when possible. For example, *a*16:0/16:0 PC represents 1-O-hexadecyl-2-palmitoyl-sn-glycero-3-phosphocholine, *e*18:0/18:2-PE represents 1-O-octadec-1′-enyl-2-octadecadienoyl-sn-glycero-3-phosphoethanolamine, 36:4 PC represents a 1,2-diacyl-sn-glycero-3-phosphocholine in which the total number of carbon from sn-1 and sn-2 is 36 and the total number of C=C double bonds from sn-1 and sn-2 is 4, and 52:3 TAG represents a TAG in which the total number of carbon from sn-1 to sn-3 is 52 and the total number of C=C double bonds is 3.

### Statistical analysis

Unless otherwise specified, all experiments were repeated three times and each biological repeat contained 2–3 technical repeats. Differences among experimental groups were determined by the unpaired Student’s *t* test (for two groups) or one-way ANOVA (for three to four groups) using Sigmaplot 13.0 (Systat Software Inc, San Jose, CA). *P* values indicating statistical significance were grouped into values of <0.05 (*), <0.01 (**) and <0.001 (***).

## RESULTS

### Targeted deletion of the endogenous *CEPT* alleles in *L. major*

*L. major* CEPT is encoded by a single copy gene on chromosome 36 (TritrypDB ID: LmjF36.5900) with well conserved, syntenic orthologs among other trypanosomatids. The predicted protein contains 417 amino acids with 6 transmembrane helices showing 30-33% identity to the CEPTs from organisms outside of kinetoplastids. To investigate the function of this enzyme in *L. major*, we first tried to generate the *CEPT*-null mutants using a classic approach based on homologous recombination (Cruz et al., 1991). With this method, we were able to replace the first *CEPT* allele with the puromycin resistance gene (*PAC*) generating *CEPT+/−* parasites (Fig. 2A). However, repeated attempts to delete the second *CEPT* allele with the blasticidin resistance gene (*BSD*) were unsuccessful, suggesting that CEPT was indispensable for promastigotes. To overcome this obstacle, we used an episome facilitated approach to acquire the chromosomal knockouts (Murta et al., 2009). Briefly, *CEPT* was cloned into the pXNG4 vector which contains genes for nourseothricin resistance (*SAT*), green fluorescent protein (GFP), and thymidine kinase (TK). The pXNG4-*CEPT* plasmid was then introduced into *CEPT+/−* to generate *CEPT+/− +*pXNG4-*CEPT*, followed by the replacement of the second chromosomal *CEPT* allele with *BSD*. With this method, parasites showing resistance to puromycin, nourseothricin, and blasticidin were readily obtained. Individual clones were isolated from these parasites by serial dilution. In a Southern blot analysis, these clones (*cept*^*−*^ *+*pXNG4-*CEPT* #2-2, 2-3 and 2-4) showed a complete loss of chromosomal *CEPT* while maintaining a high level of episomal *CEPT* (Fig. 2A). No significant difference was detected among these clones and results from *cept*^*−*^ *+*pXNG4-*CEPT* #2-3 were described in the following experiments. As shown in Fig. 2B, overexpression of *CEPT* had little impact on promastigote replication in culture. The fact that chromosomal null mutants could only be generated in the presence of a complementing plasmid suggests that CEPT is essential during the promastigote stage of *L. major.*

**Figure 2.**
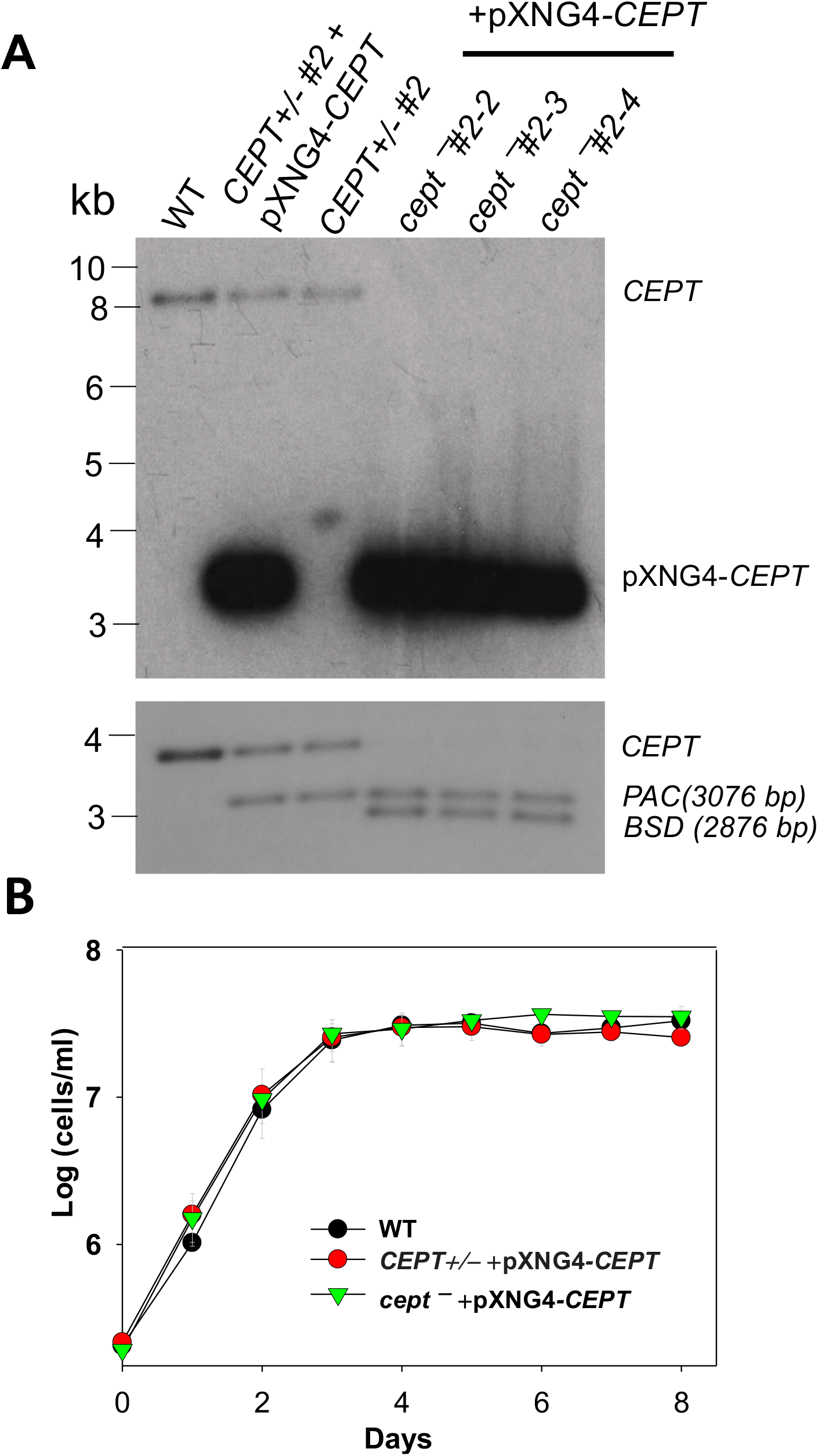
Deletion of *L. major CEPT* in the presence of pXNG4-*CEPT*. (**A**) Genomic DNA samples from WT, *CEPT* +/− clone #2, *CEPT* +/− +pXNG4-*CEPT* (clone #2), and *cept*^*−*^ *+*pXNG4-*CEPT* (clones #2-3, 2-3, and 2-4) parasites were processed for Southern blot using radiolabeled probes for the ORF (top) or flanking sequence (bottom) of *CEPT*. Bands corresponding to chromosomal *CEPT* (top: 8252 bp, bottom: 3730 bp), pXNG4-*CEPT* (top: 3475 bp), *PAC* (bottom: 3076 bp) and *BSD* (bottom: 2876 bp) were indicated. (**B**) Promastigotes of WT, *CEPT* +/− +pXNG4-*CEPT*, and *cept*^*−*^ *+*pXNG4-*CEPT* were cultivated at 27 °C in complete M199 media and culture densities were determined using a hemocytometer. Error bars represent standard deviations from three experiments.

### Expression and localization of CEPT in *L. major*

The relative transcript levels of *CEPT* in wild type (WT) promastigotes (from *in vitro* culture) and amastigotes (isolated from infected BALB/c mice) were examined by qRT-PCR. Using the 28S RNA as the internal standard, we detected a three-fold reduction in *CEPT* transcript level in amastigotes in comparison to promastigotes (Fig. 3A), suggesting that this enzyme is in lower demand during the intracellular stage. In *T. brucei*, EPT and CEPT are located in different (although overlapping) sub-compartments of the endoplasmic reticulum (ER), suggesting a spatial segregation of ether lipids (e.g. PME) and diacyl-PE/PC synthesis (Farine et al., 2015). To determine the cellular localization of CEPT in *L. major*, GFP-*CEPT* was introduced into WT promastigotes (Fig. S1). In immunofluorescence microscopy, GFP-*CEPT* exhibited a diffused and membranous pattern resembling that of the ER marker BiP (Fig. 3B-F). Quantitative analysis revealed ~73% overlap between GFP-*CEPT* and BiP (Table S1), suggesting that CEPT is primarily located at the bulk ER. This is similar to the localization of EPT and CPCT in *L. major* (Pawlowic et al., 2016; Moitra et al., 2019).

**Figure 3.**
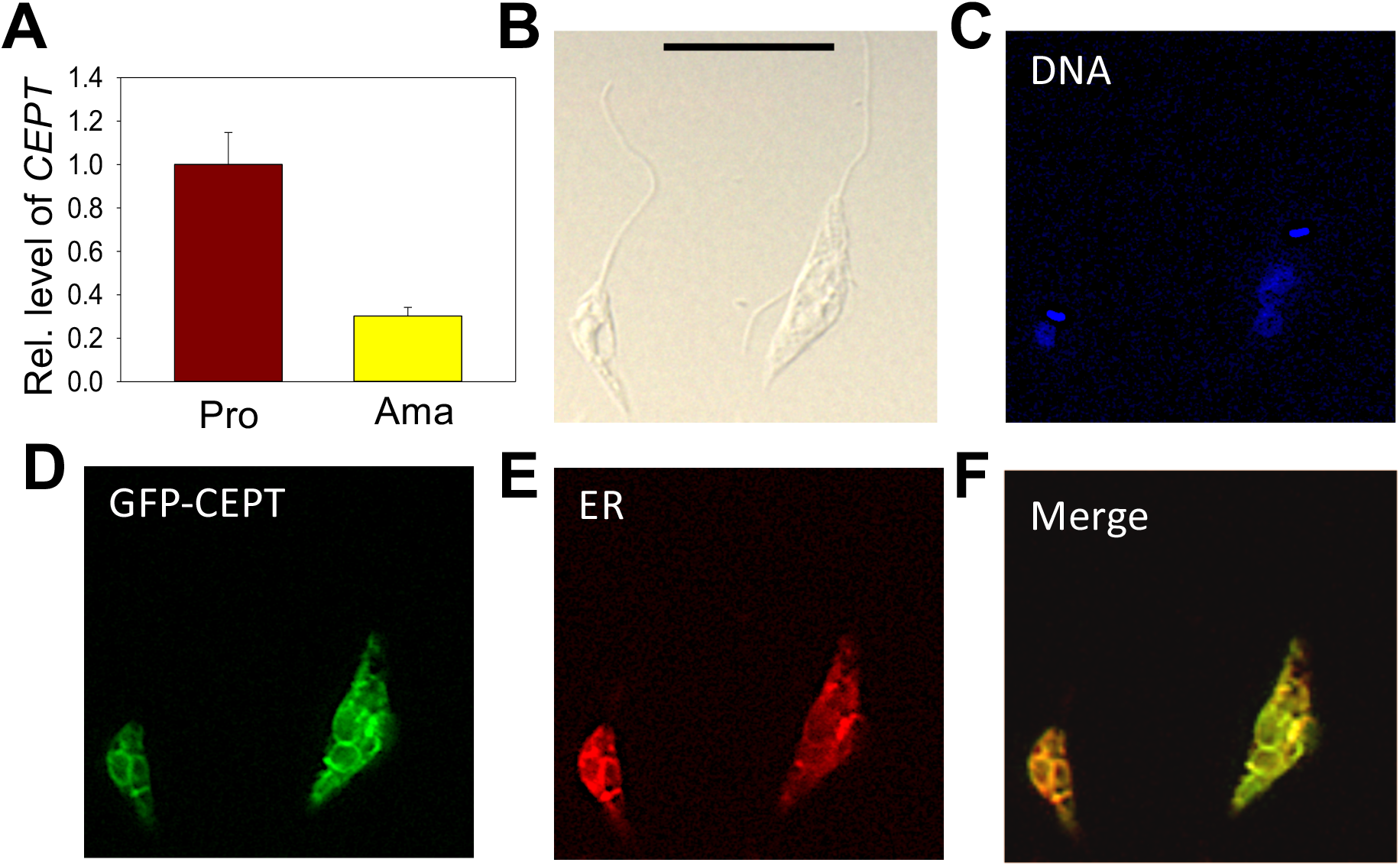
Transcript level and localization of CEPT in *L. major*. (**A**) The transcript level of *CEPT* was determined from culture promastigotes and lesion-derived amastigotes by qRT-PCR. The relative abundance of *CEPT* transcript was normalized using 28S RNA as the internal control based on ΔΔCt values. Error bars represent standard deviations from two independent biological repeats. (**B**)-(**F**) Log phase promastigotes of WT*+*pXG-*GFP-CEPT* were labeled with rabbit *anti-T. brucei* BiP antiserum followed by a goat anti-rabbit IgG-Texas Red antibody and subjected to immunofluorescence microscopy. (**B**) Differential interference contrast; (**C**) DNA staining; (**D**) GFP fluorescence; (**E**) Anti-BiP staining; (**F**) Merge of **D** and **E**. Scale bar: 10 μm. The overlap between BiP and GFP-CEPT was determined by the JaCOP Image J analysis of 136 cells (Table S1).

### CEPT is responsible for PC synthesis from CDP-choline and DAG

To determine whether *L. major* CEPT catalyzes the synthesis of PC, promastigote lysates were incubated with [^14^C]-CDP-choline and DAG for 1 hour followed by lipid extraction and thin layer chromatography (TLC). As illustrated in Fig. 4, overexpression of *CEPT* or *GFP-CEPT* led to robust production of [^14^C]-PC in the organic fraction. CEPT activity was also present in the mouse liver lysate (positive control). In comparison, this activity was barely detectable in WT parasites and completely absent in heat-inactivated lysates. Hydrolysis of CDP-choline into choline was also detected from unboiled samples. As expected, the majority of [^14^C]-CDP-choline (a substrate) was found in the aqueous phase (Fig. 4). These findings suggest that CEPT is a functional enzyme capable of generating PC from CDP-choline and DAG in *L. major.* We were unable to confirm the contribution of CEPT to PE synthesis due to the lack of commercially available, radiolabeled CDP-EtN.

**Figure 4.**
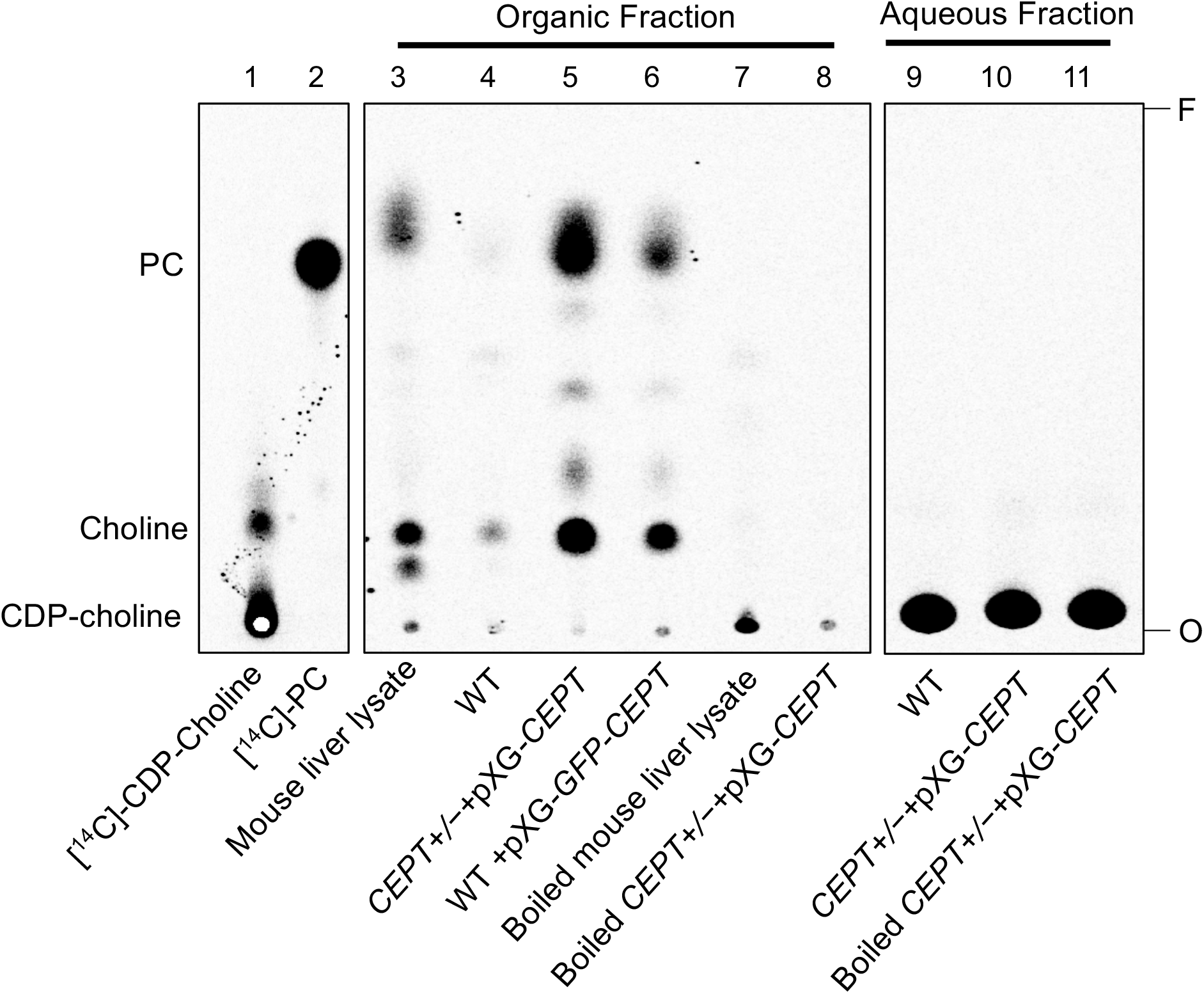
CEPT is responsible for the synthesis of PC from CDP-choline and DAG. Mouse liver lysate or promastigote lysate were incubated with [^14^C]-CDP-choline and DAG followed by extraction with 1-butanol and TLC analysis as described in *Materials and Methods*. Lane 1: [^14^C]-CDP-choline. Lane 2: [^14^C]-1,2-diarachidonyl PC. Lane 3-8: Organic fractions after 1-butanol extraction. Lane 9-11: aqueous fractions after 1-butanol extraction. O: origin. F: Solvent front. This assay was repeated three times and results from one representative experiment were shown here.

### CEPT is indispensable for *L. major* promastigotes

To critically evaluate whether CEPT is required for *L. major* promastigotes, ganciclovir (GCV) was added to *CEPT+/−* +pXNG4-*CEPT* and *cept*^*−*^ +pXNG4-*CEPT* parasites which would trigger the formation of GCV-triphosphate through the activity of TK, resulting in premature termination of DNA synthesis. If the episomal copy of CEPT in pXNG4-*CEPT* is dispensable, it would be gradually lost during *in vitro* culture in the presence of GCV (Murta et al., 2009). This process was monitored by flow cytometry (the pXNG4 plasmid contains GFP) and qPCR over multiple passages in culture (Fig. 5). As expected, when cultivated in the presence of nourseothricin (SAT) and absence of GCV, 90-95% of *CEPT+/−* +pXNG4-*CEPT* and *cept*^*−*^ +pXNG4-*CEPT* promastigotes showed high GFP expression by flow cytometry (Fig. 5A). When cultivated in the absence of GCV or nourseothricin, the *CEPT+/−+*pXNG4-*CEPT* parasites gradually lost the episome from 92% GFP-high in passage 1 to <4 % in passage 10 (Fig. 5A). In the presence of GCV and absence of nourseothricin, the reduction of GFP fluorescence was much faster in these parasites (<1% GFP-high by passage 5, Fig. 5A).

**Figure 5.**
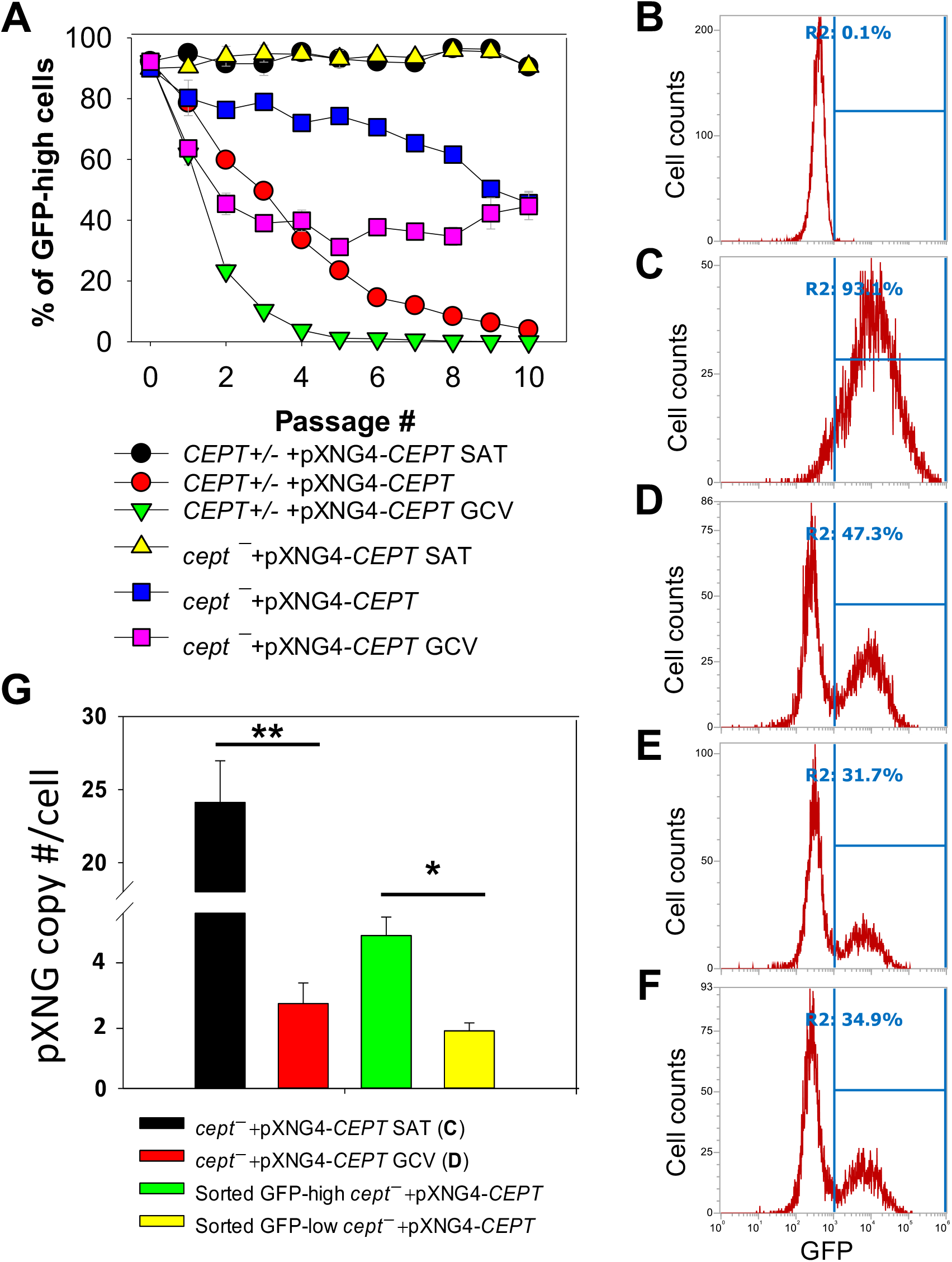
CEPT is indispensable for *L. major* promastigotes. (**A**) Promastigotes were continuously cultivated in the presence or absence of GCV or nourseothricin (SAT) and percentages of GFP-high cells were determined by flow cytometry for each passage. (**B**-**D**) After 16 passages, flow cytometry analyses were performed on WT parasites (**B**), *cept*^*−*^+pXNG4-*CEPT* parasites grown in the presence of nourseothricin (**C**) or GCV (**D**). (**E-F**) Two clones of *cept*^*−*^+pXNG4-*CEPT* were isolated from the GFP-low population in **D** by serial dilution and subjected to flow cytometry. In **B**-**F**, percentages of GFP-high cells (summarized in **A**) were indicated as R2. (**G**) The GFP-high and GFP–low parasites in **D** were separated by FACS, and plasmid copy numbers were determined by qPCR as described in *Materials and Method*s. Error bars represent standard deviations from two or three biological repeats (*: *p*<0.05, **: *p*<0.01).

Meanwhile, the *cept*^*−*^*+*pXNG4-*CEPT* parasites were 40-50% GFP-high after 10 passages in the absence of GCV (Fig. 5A). Importantly, continuous GCV treatment did not significantly reduce the GFP-high percentage in these cells (Fig. 5A-D). Thus, unlike the *CEPT+/−+*pXNG4-*CEPT* parasites which had a chromosomal *CEPT* allele, *cept*^*−*^ *+*pXNG4-*CEPT* parasites could not fully lose the episome even when they were under strong negative selection pressure.

Next, we examined if it was possible to isolate viable and replicative *CEPT*-null promastigotes without the episome. First, *cept*^*−*^*+*pXNG4-*CEPT* parasites were continuously cultivated in the presence of GCV and absence of nourseothricin for 16 passages. As shown in Fig. 5D, ~53% of these cells were classified as GFP-low and ~47% were classified as GFP-high. We then separated the GFP-low and GFP-high populations by fluorescence-activated cell sorting (FACS), followed by serial dilution to isolate clones from these populations. Importantly, when these GFP-low clones were cultivated in drug-free M199 media, 31-35% of them were GFP-high (Fig. 5E-F). Furthermore, qPCR analyses revealed averages of 1.64 copies of pXNG4-*CEPT* plasmid per cell in the sorted GFP-low clones and 4.36 copies per cell in the sorted GFP-high clones (Fig. 5G). Meanwhile, the pre-sorting population in Fig. 5D (*cept*^*−*^*+*pXNG4-*CEPT* grown in GCV for 16 passages) had 2.42 copies/cell (Fig. 5G). As a control, *cept*^*−*^*+*pXNG4-*CEPT* promastigotes cultured in the presence of nourseothricin retained 24.12 copies/cell (Fig. 5G).

Together, these results indicate that CEPT is essential for cell survival and proliferation during the promastigote stage of *L. major*.

### CEPT is dispensable for *L. major* amastigotes

To investigate whether CEPT is required during the amastigote stage, we infected BALB/c mice in the footpad with WT, *CEPT+/−* +pXNG4-*CEPT* or *cept*^*−*^*+*pXNG4-*CEPT* promastigotes. Half of the infected mice received daily treatment of GCV for 14 days and the other half received equivalent amount of sterile PBS as controls. Parasite virulence was assessed by measuring the development of footpad lesions and parasite growth in mice was determined by limiting dilution assay (LDA) and qPCR analysis. No significant difference in body weight was detected between GCV and PBS treated mice (Fig. S2). As expected, mice infected by WT parasites were not affected by GCV treatment in footpad lesion development (Fig. 6A). Similar results were observed in mice infected by *CEPT+/−* +pXNG4-*CEPT* and *cept*^*−*^*+*pXNG4-*CEPT* parasites as no significant differences were detected between the GCV- and PBS-treated groups (Fig. 6A). We then determined parasite numbers in the footpads of infected mice at 6-, 8- and 9-weeks post infection (Fig. 6B-C). While all groups showed robust parasite loads which increased over time, mice infected by *cept*^*−*^*+*pXNG4-*CEPT* (both GCV- and PBS-treated groups) had more parasites than other groups at 6- and 8-weeks post infection (Fig 6B-C, 9-week data were not available for *cept*^*−*^*+*pXNG4-*CEPT* because those mice had reached the humane endpoint after 8 weeks). These results were largely in agreement with the lesion development displayed by infected mice (Fig. 6A). Thus, *cept*^*−*^*+*pXNG4-*CEPT* parasites are fully virulent and proliferative in BALB/c mice even after GCV treatment.

**Figure 6.**
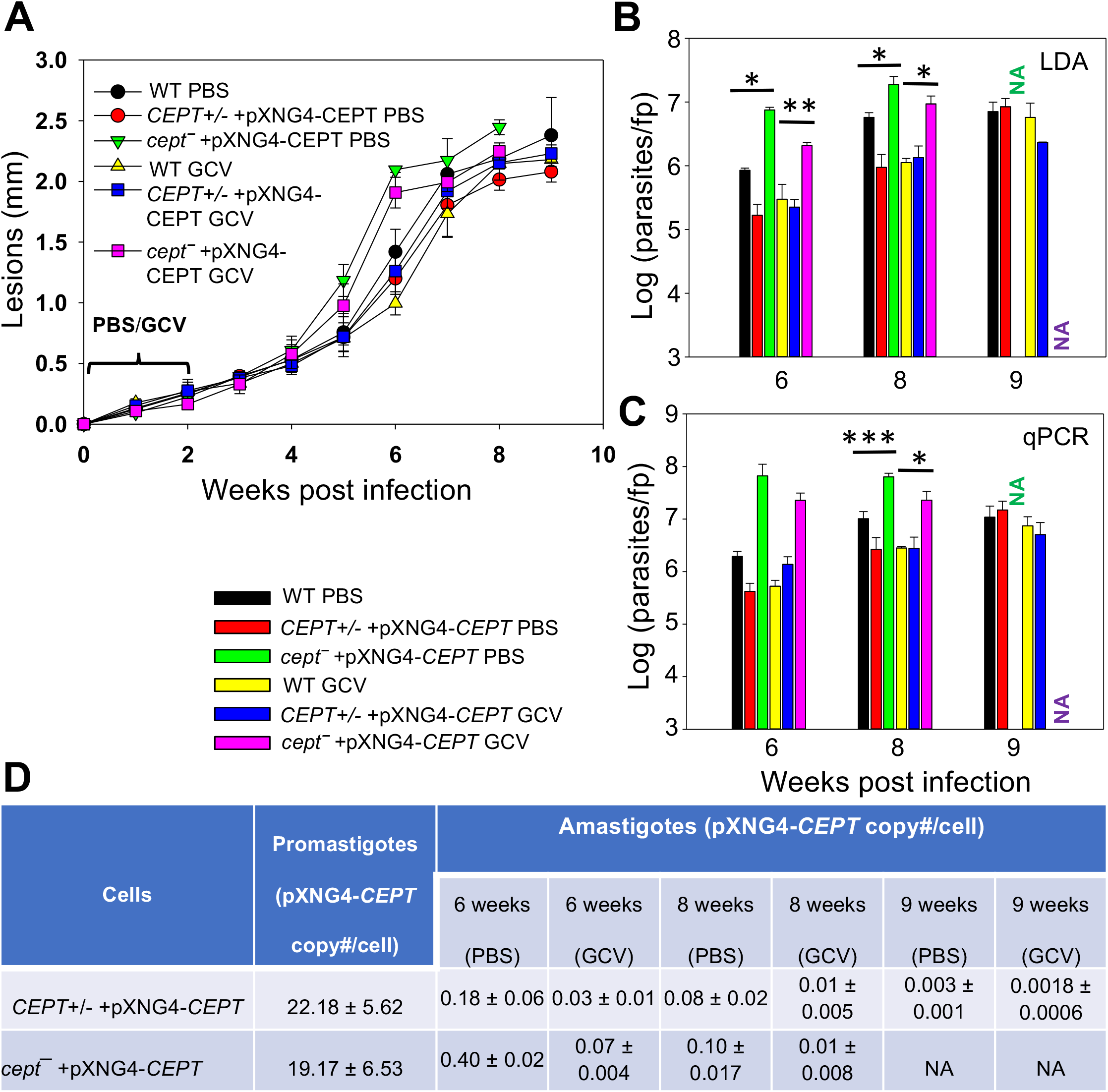
CEPT is dispensable for *L. major* amastigotes. BALB/c mice were infected in the footpads with stationary phase promastigotes and treated with either GCV or PBS from day 1 to day 14 post infection as described in *Materials and Methods*. (**A**) Footpad lesions were recorded weekly. (**B**)-(**C**) Parasite numbers in the infected footpads were determined at the indicated times by limiting dilution assay (**B**) and qPCR (**C**). (**D**) The pXNG4-*CEPT* copy numbers in promastigotes and amastigotes (#/cell ± SDs) were determined by qPCR. The 9 weeks data for *cept*− +pXNG4-*CEPT* amastigotes were not available (NA) because the infected mice had reached the humane endpoint at 8 weeks. Error bars represent standard deviations (*: *p* < 0.05, **: *p* < 0.01, ***: *p* < 0.001).

Next, we examined whether *CEPT+/−* +pXNG4-*CEPT* and *cept*^*−*^*+*pXNG4-*CEPT* parasites would retain the pXNG4-*CEPT* plasmid in mice. To do so, qPCR analyses were performed on the amastigote DNA isolated from infected mice at 6-, 8-, and 9-weeks post infection using primers for the pXNG4 plasmid (to determine plasmid copy number) and for the *L. major* 28S rRNA gene (to determine parasite number). The average plasmid copy number per cell was determined by dividing the total number of pXNG4 plasmid molecules with the total number of parasites. As illustrated in Fig. 6D, *CEPT+/−* +pXNG4-*CEPT* amastigotes contained 0.08-0.18 copies of the episome per cell in PBS-treated mice at weeks 6-8 post infection, and GCV-treatment further reduced the average copy number to 0.01-0.03 per cell. At week 9 post infection, the pXNG4 copy number became nearly undetectable (<0.01, Fig. 6D). Importantly, similar results were obtained for the *cept*^*−*^*+*pXNG4-*CEPT* amastigotes as the episome copy numbers were 0.10−0.40 copies per cell in PBS-treated mice and 0.01−0.07 copies per cell in GCV-treated mice (Fig. 6D). As controls, promastigotes of *CEPT+/−* +pXNG4-*CEPT* and *cept*^*−*^*/+*pXNG4-*CEPT* cultivated in the presence of nourseothricin contained 19-24 copies of pXNG4-*CEPT* per cell, and those cultured in the presence of GCV contained 1.6-4.4 copies per cell (Fig. 6D and Fig. 5G). The fact that *cept*^*−*^*+*pXNG4-*CEPT* amastigotes were fully virulent and replicative with very few copies of the episome (much less than one plasmid per cell) suggests that CEPT is dispensable during the mammalian (intracellular) phase. These findings are distinct from our previously reported *fpps−+*pXNG4-*FPPS* parasites (Mukherjee et al., 2019). Thus, the *de novo* synthesis of PC is only required for the promastigote but not amastigote stage in *L major*.

Our conclusion was also supported by comparing the parasite loads data from LDA and qPCR (Fig. 6B-C). For WT and *CEPT+/−* +pXNG4-*CEPT* parasites, results from LDA and qPCR were largely comparable (Fig. 6B-C). In contrast, for *cept*^*−*^*+*pXNG4-*CEPT*, the qPCR results were 4-10 times higher than the corresponding LDA results (2-6 × 10^7^ vs 2-20 × 10^6^, Fig. 6C). While qPCR revealed the total number of amastigotes in the infected footpad, LDA only detected those promastigotes that had successfully converted from amastigotes and replicated *in vitro*. It appears that only a fraction of *cept*^*−*^*+*pXNG4-*CEPT* amastigotes can be converted into proliferative promastigotes due to their low episome copy number (more on this point below).

### *Cept*^*−*^*+* pXNG4-*CEPT* parasites regain the plasmid after conversion from amastigotes to promastigotes

The apparent lack of episome in *cept*^*−*^*+*pXNG4-*CEPT* amastigotes (Fig. 6D) prompted us to examine their ability to convert to promastigotes and proliferate in culture in more detail. First, *CEPT+/−* +pXNG4-*CEPT* and *cept*^*−*^*+*pXNG4-*CEPT* amastigotes were isolated from infected mice (Fig. 6) and allowed to recover in complete M199 media in the absence or presence of nourseothricin (“no drug” or “+ SAT” in Fig. 7A). After amastigotes had converted into promastigotes (in 2-4 days), they were cultivated in the absence or presence of nourseothricin for 10 consecutive passages and their GFP fluorescence levels were determined by flow cytometry (Fig. 7B-E). Without the positive selection drug nourseothricin, *cept*^*−*^*+*pXNG4-*CEPT* parasites isolated from PBS-treated mice showed 40-60% GFP-high cells, whereas *CEPT+/−* +pXNG4-*CEPT* parasites lost most of the GFP-high cells (Fig. 7B). Very similar results were observed for the parasites isolated from GCV-treated mice except that the GFP-high population in *cept*^*−*^*+*pXNG4-*CEPT* was ~15% in the first passage before going up to 50-60% in the later passages (Fig. 7C). The slower recovery may be due to the extremely low episome copy number in parasites isolated from in GCV-treated mice (0.01-0.07/cell for *cept*^*−*^*+*pXNG4-*CEPT*, Fig. 6D).

**Figure 7.**
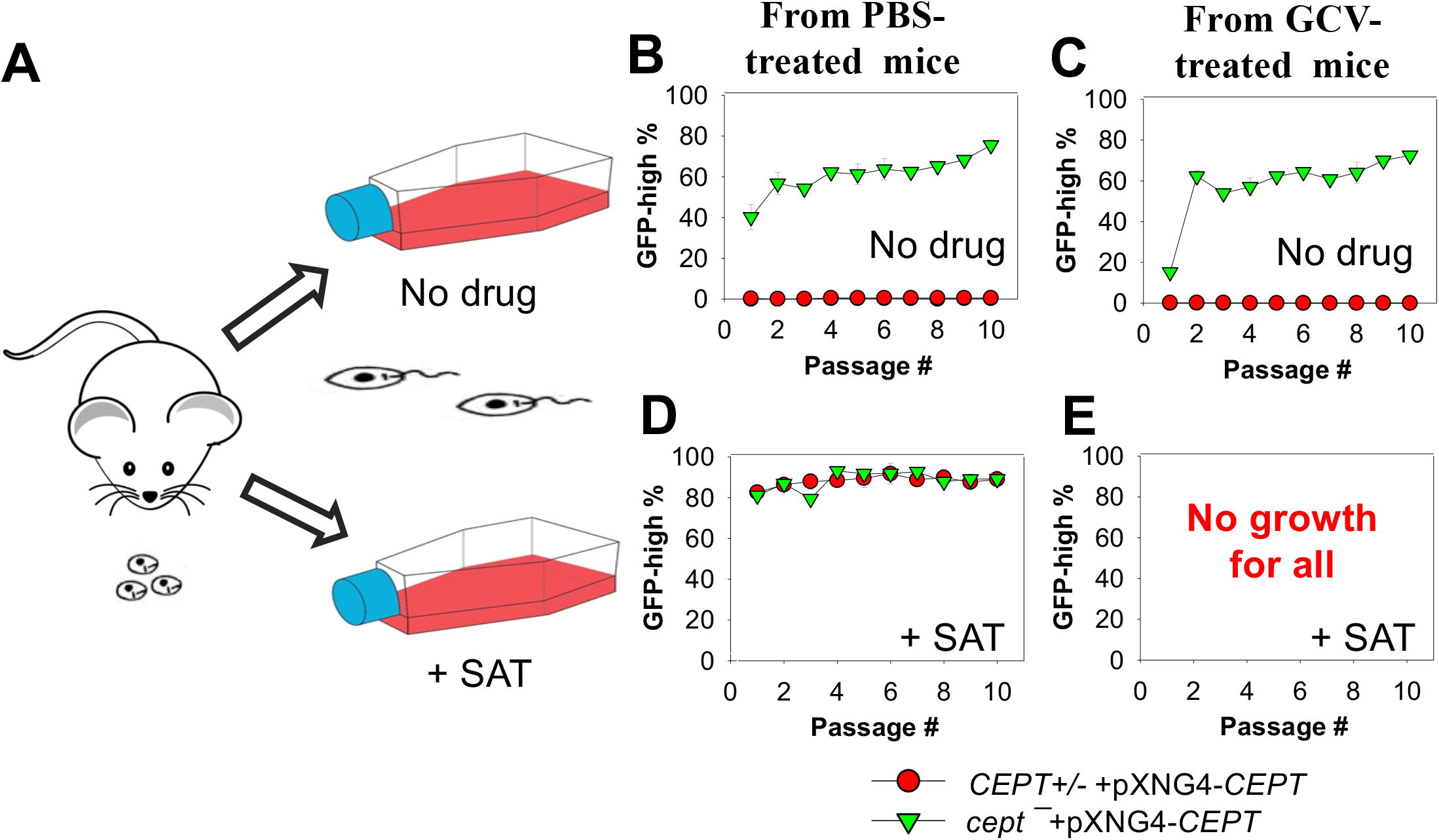
*Cept*^*−*^+pXNG4-*CEPT* parasites restore their episome levels after converting to promastigotes. (**A**) Amastigotes from infected BALB/c mice were allowed to recover in complete M199 media in the absence or presence of 200 μg/ml of nourseothricin (+ SAT). Promastigotes were then continuously cultivated in the absence (**B** and **C**) or presence (**D** and **E**) of nourseothricin for 10 passages and percentages of GFP-high cells were analyzed by flow cytometry for each passage. It is of note that no cells from GCV-treated mice were able to recover in nourseothricin (**E**). For **B**-**D**, error bars represent standard deviations from two independent biological experiments (two technical duplicates each time).

Meanwhile, if parasites were continuously cultivated in the presence of nourseothricin, both *cept*^*−*^*+*pXNG4-*CEPT* and *CEPT+/−* +pXNG4-*CEPT* exhibited 80-90% GFP-high cells if they were isolated from PBS-treated mice (Fig. 7D). However, parasites isolated from GCV-treated mice failed to grow in the presence of nourseothricin (Fig. 7E). These results suggest that the extremely low episome levels in *CEPT+/−* +pXNG4-*CEPT* and *cept*^*−*^*+*pXNG4-*CEPT* amastigotes isolated from GCV-treated mice (0.01−0.07 copies per cell, Fig. 6D) prevented these parasites from converting into proliferative promastigotes in the presence of nourseothricin.

In addition, the effect of GCV on promastigote proliferation was examined (Fig. S3). If *cept − +*pXNG4-*CEPT* amastigotes were allowed to convert into promastigotes in the absence of GCV (“no drug” in Fig. S3A), no significant difference in replication was observed (Fig. S3B). This was consistent with the rapid restoration of episome level in *cept− +*pXNG4-*CEPT* (Fig. 7). However, if these amastigotes were forced to convert into promastigotes in the presence of GCV (“+ GCV” in Fig. S3A), they showed a 24-96 hours delay during the first passage (“P1” in Fig. S3B). This delay was likely caused by the difficulty of these cells trying to increase their episome copy numbers to support growth while balancing the toxic effect from GCV. They were able to proliferate normally after the first passage (“P2-P4” in Fig. S3B), suggesting adaptation to limiting the pXNG4 copy number to proper levels that satisfy the need of PC synthesis while limiting toxicity. We also analyzed the pXNG4-*CEPT* DNA from those recovered culture promastigotes (Fig. 7 and Fig. S3) and did not detect any mutations in the plasmid based on restriction enzyme digestion and sequencing.

These findings suggest that when *cept*^*−*^ +pXNG4-*CEPT* amastigotes were allowed to recover during the initial passage in culture, they quickly elevated the plasmid copy numbers while transitioning to promastigotes. This occurred even in the absence of nourseothricin (the selection drug for pXNG4 plasmid) and presence of GCV due to the essentiality of CEPT in promastigotes. In contrast, for *CEPT+/−* +pXNG4-*CEPT*, the plasmid copy number did not increase without strong positive selection.

### *Cept*^*−*^*+* pXNG4-*CEPT* parasites showed similar PC and PE contents as WT parasites

Next, we determined the lipid composition in *cept − +*pXNG4-*CEPT* parasites to evaluate the impact of CEPT on PC/PE production. First, total lipids from culture promastigotes were examined by ESI-MS in the positive ion mode to detect PC (as protonated [M+H]^+^ ions) and negative ion mode to detect PE (as deprotonated [M-H]^-^ ions). As indicated in Fig. S4A, the majority of PC in promastigotes were 1,2-diacyl types with polyunsaturated FAs at the sn-2 positions (e.g., 18:2/22:6 PC, 18:3/22:6 PC and 16:1/20:4 PC) while ether PC (e.g., a18:0/20:2 PC and e18:1/24:6 PC) were less common. These findings were largely in agreement with previous reports (Weingartner et al., 2012; Williams et al., 2012). No statistically significant difference in PC composition was detected between WT and *cept*^*−*^*+*pXNG4-*CEPT* promastigotes (Fig. S4A). Similarly, the composition for PE (mainly e18:0/18:2 PE and e18:0/18:1 PE, both are PME) in *cept*^*−*^*+*pXNG4-*CEPT* were very similar to WT promastigotes (Fig. S4B) (Zhang et al., 2007).

Finally, we examined the lipids from lesion-derived amastigotes (Fig. S5) and uninfected mouse footpads. As summarized in Fig. 8A, the PC in WT amastigotes included both ether lipids (e.g., a16:0/16:0 PC, a16:0/18:2 PC, and a16:0/18:1 PC) and 1,2-diacyl types (e.g., 36:4 PC, 36:2 PC and 34:2 PC). Importantly, very similar PC composition was found in *cept*^*−*^*+*pXNG4-*CEPT* amastigotes despite the near complete loss of *CEPT* gene in these parasites (Fig. 8A and 6D). Thus, *L. major* amastigotes can acquire the majority of their PC in the absence of *de novo* synthesis. While some of these amastigote PC (e.g., 36:4 PC, 34:2 PC and 34:1 PC) were also found in uninfected mouse tissue, the majority of amastigote PC were of low abundance or undetectable in the host (Fig. 8A) (e.g., a16:0/16:0 PC, a16:0/18:2 PC and a16:0/18:1 PC). These findings suggest that amastigotes can generate PC through the uptake and remodeling of host PC.

**Figure 8.**
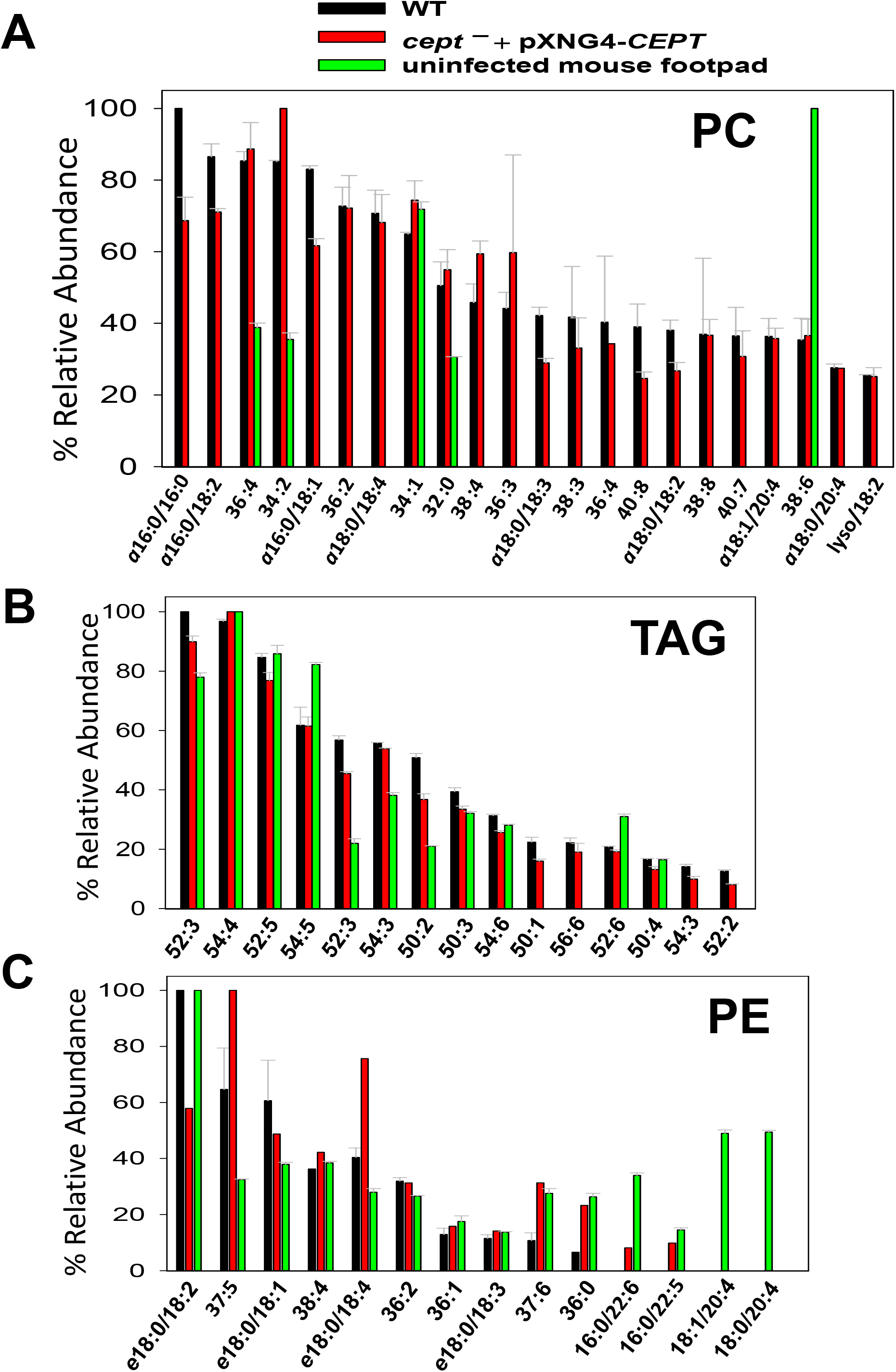
*Cept*^*−*^+pXNG4-*CEPT* amastigotes show similar composition of PC and PE as WT amastigotes. Amastigotes were isolated from the footpad of infected BALB/c mice. Lipids from partially purified amastigotes and uninfected mouse footpad tissue were analyzed by ESI-MS in the positive ion mode (**A** and **B**) and negative ion mode (**C**). Compositions for PC (**A**), TAG (**B**), and PE (**C**) were summarized. Only major lipid species (>20% for PC and >10% for TAG and PE) by relative abundance are shown. Predicted formulas and molecular weights were indicated for each lipid species (*a*: 1-alkyl phospholipids, *e*: 1-alkenyl phospholipids, lyso: 1-lysophospholipids).

In comparison to PC, we found higher degrees of overlap in the composition of triacylglycerol (TAG) and PE between amastigotes and mouse tissue (Fig. 8B and C). Again, no major differences in TAG and PE composition were detected between WT and *cept*^*−*^*+*pXNG4-*CEPT* amastigotes (Fig. 8B and C). In summary, our data indicate that the *de novo* PC synthesis is dispensable for the survival and proliferation of intracellular amastigotes which can salvage and remodel lipids from the host.

## DISCUSSION

*Leishmania* parasites can synthesize PC from multiple routes including the choline branch of the Kennedy pathway, the N-methylation of PE and the salvage/remodeling of host lipids (Fig. 1). In this study, we characterized the gene encoding CEPT in *L. major* which catalyzes the formation of PC from CDP-choline and DAG, the final step of *de novo* PC synthesis (Fig. 1 and 4). Similar to CPCT and EPT, CEPT is primarily located at the ER (Fig. 3) (Pawlowic et al., 2016; Moitra et al., 2019). While both the choline branch and ethanolamine branch are essential for survival in *T. brucei* (Gibellini et al., 2009; Gibellini and Smith, 2010), *Leishmania* parasites can tolerate the loss of several Kennedy pathway enzymes presumably from the compensatory effects of other pathways, as evidenced by the *cpct*^*−*^ and *ept*^*−*^ mutants (Pawlowic et al., 2016; Moitra et al., 2019). However, we could not generate the chromosomal *CEPT* null mutant in culture without first introducing a complementing episome (Fig. 2). Attempts to eliminate the GFP-containing pXNG4-*CEPT* episome via prolonged GCV-treatment in culture were unsuccessful in *cept*^*−*^+pXNG4-*CEPT*, as 40-50% of parasites remained GFP-high (Fig. 5A). Even after cell sorting and serial dilution, clones derived from the GFP-low population still displayed 31-35% GFP-high with 1.5-2 copies of pXNG4-*CEPT* episome per cell (Fig. 5D-G). Thus, those GFP-low *cept*^*−*^+pXNG4-*CEPT* parasites likely contained the pXNG4-*CEPT* episome at a low but detectable level or they could not proliferate without the GFP-high cells. Interestingly, *Leishmania* and *T. brucei* parasites are known to produce and exchange extracellular vesicles among individual cells (Douanne et al., 2020). Such transfer of cellular material may contribute to the survival of *cept*^*−*^+pXNG4-*CEPT* parasites after GCV treatment.

Because CEPT is also expected to catalyze the synthesis of diacyl-PE, its deletion would not only block the choline branch of the Kennedy pathway, but also reduce PC synthesis from PE N-methylation (Fig. 1). Therefore, while *L. major* promastigotes can withstand the deletion of CPCT, a complete loss of CEPT would drastically compromise their ability to generate PC. The essentiality of *CEPT* in promastigotes suggests that the salvage pathway by itself cannot produce enough PC (the most abundant lipids) to support parasite proliferation in culture.

In contrast to the promastigote stage, CEPT appears to be dispensable during the intracellular amastigote stage. In mice infected by *cept*^*−*^+pXNG4-*CEPT* parasites, GCV treatment was able to dramatically reduce the episome level to 0.01-0.07 copy per cell after 6-8 weeks (Fig. 6D). While this analysis was done at the DNA level, it suggests that the majority of *cept*^*−*^ +pXNG4-*CEPT* amastigotes lack CEPT mRNA and protein. Nonetheless, these chromosomal-null mutants were fully virulent and replicative in mice (Fig. 6A-C). In fact, these *cept*^*−*^+pXNG4-*CEPT* parasites showed slightly better growth than WT parasites in mice, suggesting that they gained a small advantage by losing *de novo* PC synthesis (Fig. 6B-C). Additionally, lipidomic analysis revealed very similar PC contents between WT and *cept*^*−*^+pXNG4-*CEPT* amastigotes (Fig. 8A). The total PC levels in WT and *cept*^*−*^+pXNG4-*CEPT* amastigotes also appeared to be similar based on the signal intensity from mass spectrometry. Thus, *L. major* amastigotes are capable of generating sufficient amounts of PC in the absence of *de novo* synthesis. This is consistent with the three-fold reduction of *CEPT* transcript level when parasites transition from promastigotes to amastigotes (Fig. 3A).

In theory, amastigotes could acquire PC from the mammalian host, either by directly incorporating host lipids or converting them into their own. As indicated in Fig. 8A, while certain PC types were present in both amastigotes and mice, the majority of amastigote PC including the most abundant species like a16:0/16:0 PC and a16:0/18:2 PC were of low abundance or undetectable in uninfected mouse tissue, suggesting that they were modified from host lipids. The mechanism by which amastigotes remodel host lipids is under investigation. In many eukaryotes, phospholipase A2 (PLA2) removes fatty acids at the sn-2 position of PC, and the resulting lysophosphatidylcholine can be re-acylated at the sn-2 position to yield a different PC (Lands cycle)(Lands, 1958). Genes encoding PLA2 and lysophosphatidylcholine acyltransferase are present in *Leishmania* genomes suggesting that they can carry out lipid remodeling via deacylation/re-acylation. The Lands cycle is also present in the intestinal protozoan *Giardia lamblia* which has a limited capacity for *de novo* lipid synthesis (Das et al., 2001). In addition to PC remodeling and *de novo* synthesis, it is also possible for amastigotes to generate PC through the N-methylation of salvaged PE (Fig. 1).

Comparing to PC, higher degrees of overlap were observed in the compositions of triacylglycerol (TAG) and PE between amastigotes and uninfected mouse tissue (Fig. 8B-C). These findings support the direct incorporation of these lipids by amastigotes similar to the reported uptake of host TAG by intracellular *T. cruzi* parasites (Gazos-Lopes et al., 2017), although further investigation is warranted. Overall, the ability of *Leishmania* amastigotes to take up and remodel host PC is in agreement with their propensity to incorporate host sphingolipids and cholesterol (Zhang et al., 2005; Xu et al., 2014). Since *Leishmania* amastigotes spend the majority of their time in mammals in a slow growing, quiescent state (estimated doubling time: 60 hours) (Saunders et al., 2014; Mandell and Beverley, 2017), the salvage pathway (which is less energy intensive than *de novo* synthesis) seems to fit the intracellular stage. In contrast, the *de novo* synthesis is required to generate sufficient amount of lipids to support parasite replication during the promastigote stage (doubling time: 6-8 hours). The ability of *Leishmania* parasites to acquire sufficient lipids via either *de novo* synthesis or salvage/remodeling (the metabolic flexibility) may allow them to adapt to a diverse range of host cells and host species. Future studies will elucidate how amastigotes regulate the uptake and remodeling of different classes of lipids and explore the potential of blocking the transfer of host lipids to parasites as a novel therapeutic.

## ACKNOWLEDGEMENTS

We thank Dr. Jay Bangs (University at Buffalo, SUNY) for providing the rabbit anti-*T. brucei* BiP antiserum and Veronica Hernandez (Texas Tech University) for technical assistance. This work was supported by the US National Institutes of Health (AI099380 for KZ and P41-GM103422, P60-DK20579, P30-DK56341 for the Biomedical Mass Spectrometry Resource at Washington University in St. Louis, MO, USA).

