## Supplemental data for "*De novo* synthesis of phosphatidylcholine is essential for the promastigote but not amastigote stage in *Leishmania major*"

**Table S1. Quantitative analysis of colocalization between GFP-CEPT and the ER marker BiP**

| GFP-CEPT Image # | # of Cells analyzed | Pearson's Coefficient |
|---|---|---|
| 1 | 4 | 0.809 |
| 2 | 7 | 0.754 |
| 3 | 5 | 0.755 |
| 4 | 3 | 0.68 |
| 5 | 9 | 0.742 |
| 6 | 3 | 0.713 |
| 7 | 6 | 0.756 |
| 8 | 23 | 0.796 |
| 9 | 15 | 0.734 |
| 10 | 8 | 0.698 |
| 11 | 14 | 0.773 |
| 12 | 10 | 0.737 |
| 13 | 9 | 0.783 |
| 14 | 9 | 0.761 |
| 15 | 6 | 0.736 |
| 16 | 5 | 0.519 |

Total cells analyzed: 136. Average Pearson's Coefficent ± SD: 0.734 ± 0.04

**Table S2. List of oligonucleotides used in this study**

| Primer # | Primer name | Sequence |
|---|---|---|
| #129 | 5' *CEPT* ORF | GATCAGGGATCCACCATGCCCCCGAAGTCGATGGC |
| #130 | 3' *CEPT* ORF | GATCATGGATCCCTAATCTGACTTATTCGGTT |
| #137 | *CEPT* 5' UTR Upstream | GATCATGAATTCGACGGAACTCTTAGCCACTC |
| #138 | *CEPT* 5' UTR Downstream | GTCAGCGGATCCGATCTAACTAGTCTTCGTTCTCCTTGTTTTGG |
| #139 | *CEPT* 3' UTR Upstream | GATCATGGATCCAGCAGCTGCGAAGTCCGCCC |
| #140 | *CEPT* 3' UTR Downstream | GATCATAAGCTTTACACCACCTCCTCGTCAAG |
| #784 | C*EPT* qRT forward | GGAGAGTATCAACCCGCTCG |
| #785 | *CEPT* qRT reverse | CAGTACCGCTGCAGGACATA |
| #780 | 28S rRNA gene forward | AAGATGGACCGGCCTCTAGT |
| #781 | 28S rRNA gene reverse | ATCCTTCCCCGCTCCAGTAT |
| #782 | pXNG4 forward | CCCGACAACCACTACCTGAG |
| #783 | pXNG4 reverse | GTCCATGCCGAGAGTGATCC |

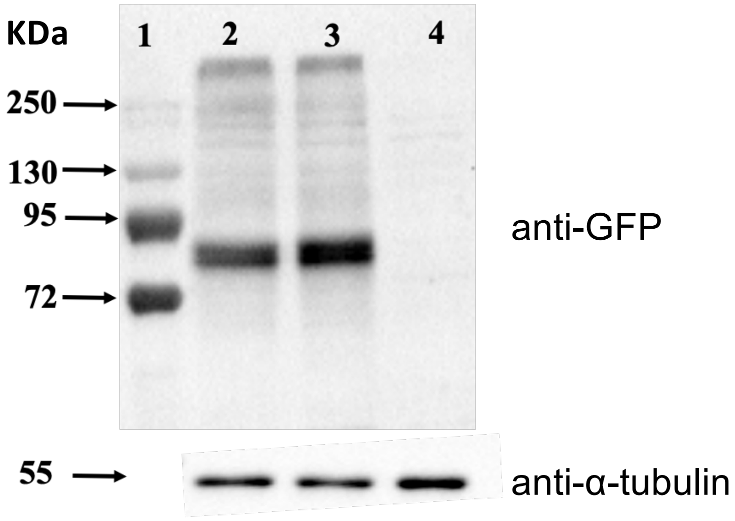


**Figure S1.** Validation of GFP-tagged CEPT expression by Western blot. Cell lysates from log phase *CEPT*+/−+pXG-*GFP-CEPT* clone 1 and 2 (Lane 2 and 3 respectively) and LV39WT promastigotes (lane 4) were analyzed by Western blot using antibodies against GFP (top) or α-tubulin (bottom). Lane 1: molecular weight marker.

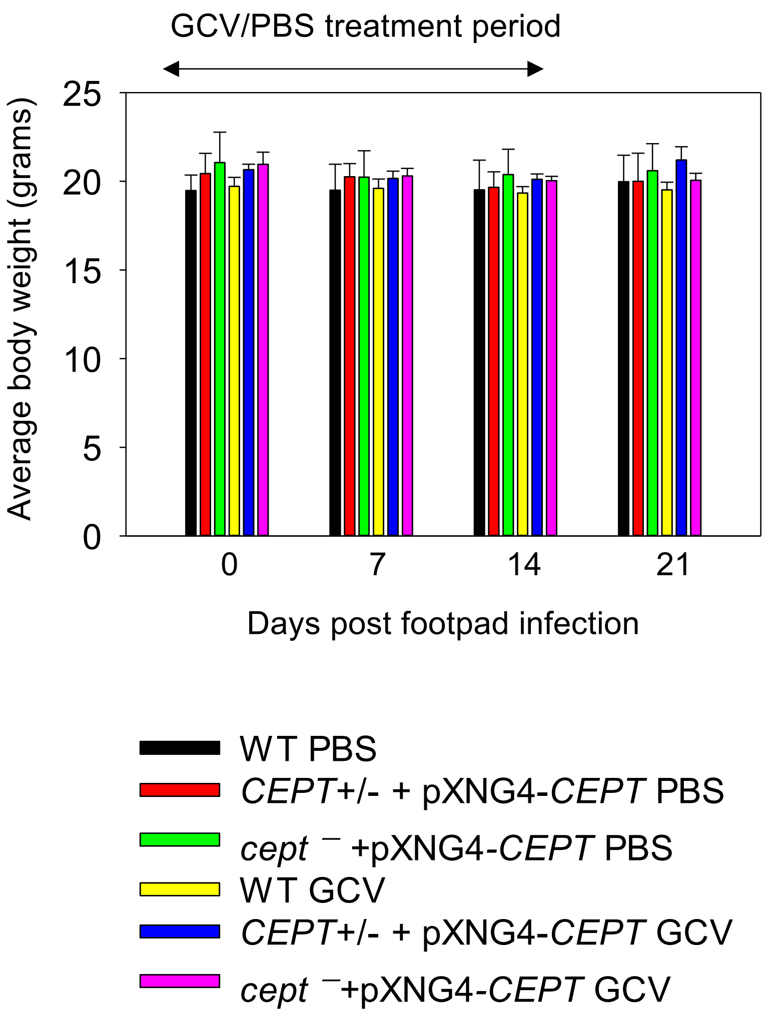


**Figure S2. GCV treatment did not affect the body weights of infected BALB/c mice.** Following infection with stationary phase promastigotes, half of the mice were treated daily with GCV for 14 days and the other group were treated with PBS. Mouse body weights were measured once a week for 3 weeks post infection. Error bars represent standard deviations (5 mice per group).

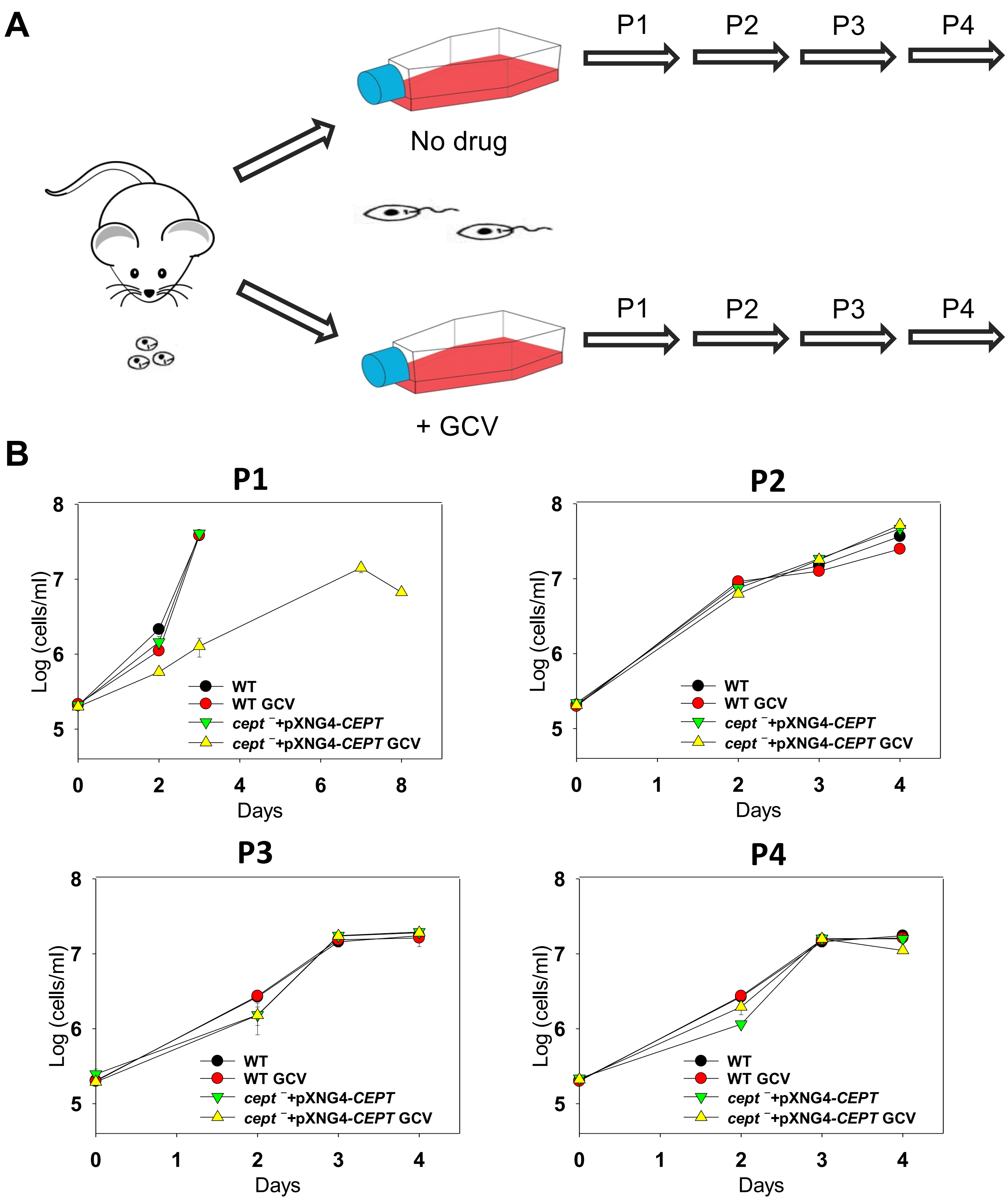


**Figure S3. GCV treatment delayed the growth of *cept* ⁻+pXNG4-*CEPT* promastigotes in the initial passage after recovery from mice.** (**A**) Experimental scheme. Amastigotes were isolated from mice and allowed to recover in complete M199 media in the absence or presence 50 µg/ml of GCV. (**B**) In subsequent passages (P1-P4), promastigotes were inoculated at 2.0 x $10^5$ cells/ml in the absence or presence 50 µg/ml of GCV. For each passage, culture densities were determined daily.

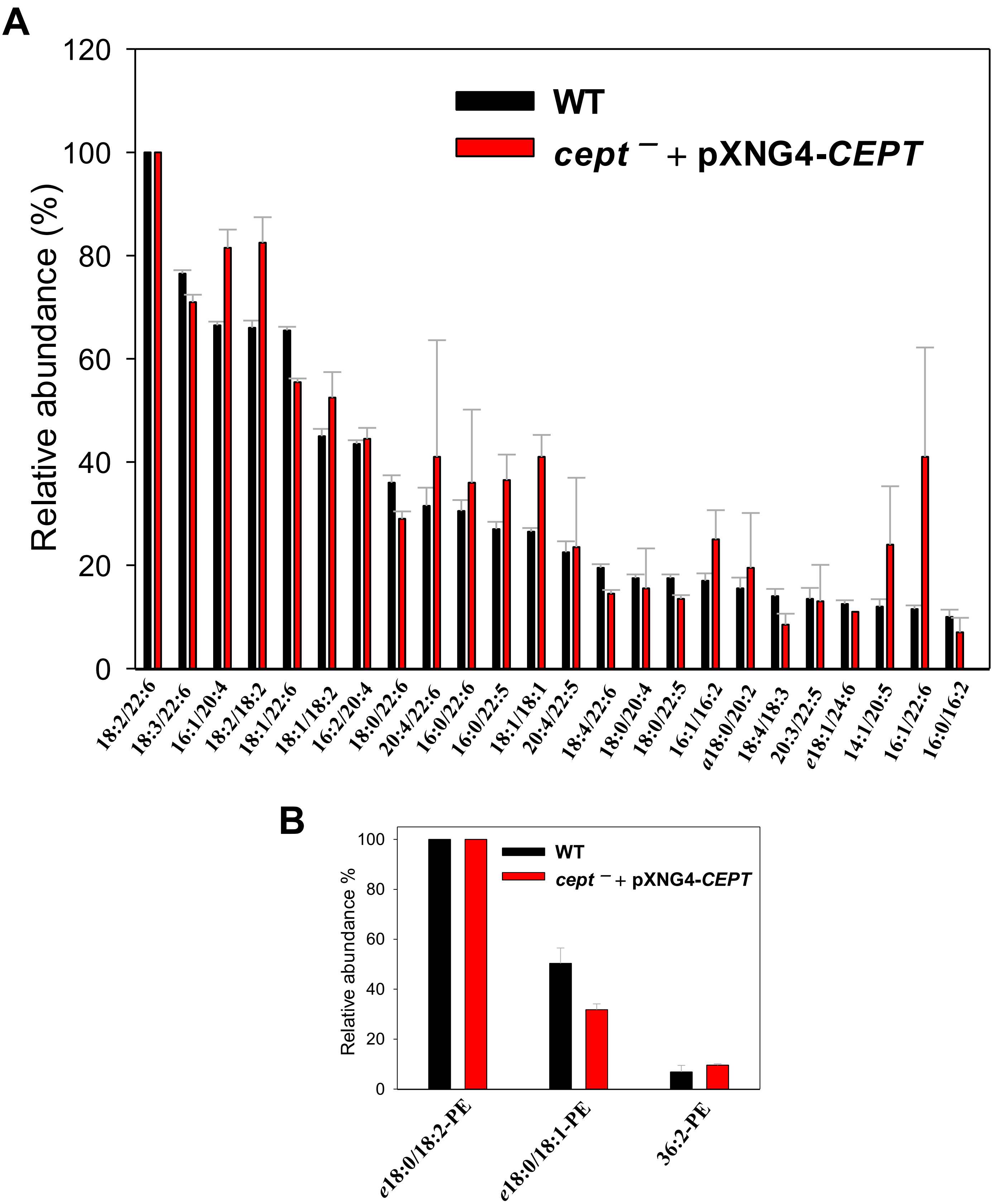


**Figure S4. *Cept* $^{-}$+pXNG4-*CEPT* promastigotes show similar phospholipid composition as WT promastigotes.** Lipids from log phase promastigotes were analyzed by ESI-MS in the positive ion mode (**A**: for PC) and negative ion mode (**B**: for PE). Predicted fatty acyl constituents were indicated. Only major PC and PE species (>5% by relative abundance) are shown.

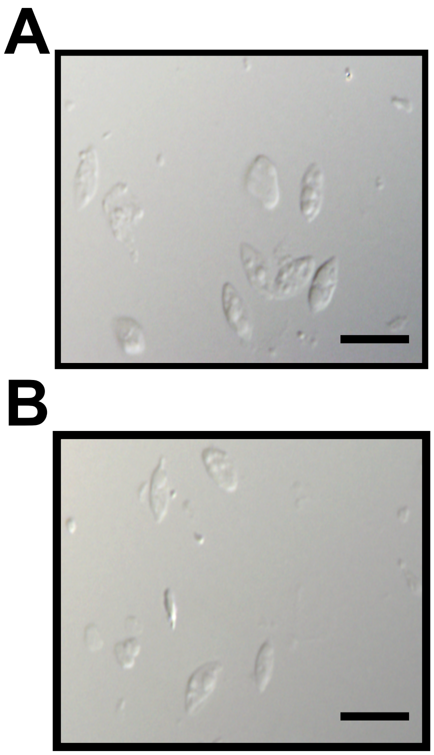


**Figure S5. DIC images of partially purified amastigotes.** Amastigotes of WT (**A**) and *cept*$^{-}$+pXNG4-*CEPT* (**B**) were isolated from infected BALB/c mice and partially purified as described in Materials and Methods. Scale bars: 10 μm.
